# EPAC1 regulates endothelial Annexin A2 cell surface translocation and plasminogen activation

**DOI:** 10.1101/187559

**Authors:** Wenli Yang, Fang C. Mei, Xiaodong Cheng

## Abstract

Annexins, a family of highly conserved calcium-and phospholipid-binding proteins, play important roles in a wide range of physiological functions. Among the twelve known annexins in human, Annexin A2 (AnxA2) is one of the most extensively studied and has been implicated in various human diseases. AnxA2 can exist as a monomer or a heterotetrameric complex with S100A10 (P11) and plays a critical role in many cellular processes including exocytosis/endocytosis and membrane organization. At the endothelial cell surface, (AnxA2•P11)_2_ tetramer, acting as a coreceptor for plasminogen and tissue plasminogen activator (t-PA), accelerates t-PA dependent activation of the fibrinolytic protease, plasmin, the enzyme responsible for thrombus dissolution and degradation of fibrin. This study shows that exchange proteins directly activated by cAMP isoform 1 (EPAC1) interacts with AnxA2 and regulates its biological functions by modulating its membrane translocation in endothelial cells. Using genetic and pharmacological approaches, it is demonstrated that EPAC1, acting through the PLCε-PKC pathway, inhibits AnxA2 surface translocation and plasminogen activation. These results suggest that EPAC1 plays a role in the regulation of fibrinolysis in endothelial cells and may represent a novel therapeutic target for disorders of fibrinolysis.

## INTRODUCTION

The plasminogen-plasmin system is involved in multiple physiological and pathological processes including fibrinolysis, angiogenesis, embryo implantation, fibrosis and wound healing, as well as cancer metastasis and invasion (*1*). Proper regulation of plasminogen and plasmin conversion on the vascular endothelial cell surface is critical for maintaining the fibrinolytic balance of blood vessel. The heterotetrameric (AnxA2•P11)_2_ complex on the cell surface promotes the catalytic efficiency of tissue plasminogen activator (tPA)-dependent plasminogen activation by serving as an assembly site for plasminogen and tPA (*2*). While plasmin activation induced by cell surface (AnxA2•P11)_2_ complex is less potent than that is induced by the fibrin during classic fibrinolysis, it may represent an important surveillance mechanism for removing minor fibrin resulted from day-to-day vascular injury or stress (*3*).

AnxA2 is a 36 KD protein containing an N-terminal hydrophilic tail and a core domain of four helical, Ca^2+^-binding “annexin” repeats, which are important for Ca^2+^-dependent membrane binding. While AnxA2 can exist as a monomer in cytosol, it can also form a heterotetrameric complex with P11, also known as S100A10. Binding of P11 enhances AnxA2’s affinity to Ca^2+^ and is required for the translocation and presentation of AnxA2 to outer leaflet of the plasma membrane (*4*). In the absence of Annexin, P11 is polyubiquitinated and rapidly degraded in a proteasome-dependent manner. Interaction between the N-terminal tail of AnxA2 and P11 stabilizes P11 by masking the polyubiquitination signal on P11 (*5*). This modulatory mechanism ensures that the translocation of AnxA2 is a highly dynamic process and precisely attuned to the intracellular level of P11.

The regulation of endothelial AnxA2 cell surface translocation is incompletely understood. Extensive studies have revealed that the translocation of the (AnxA2•P11)_2_ complex is tightly regulated in response to various physiological stimuli or stress signals such as thrombin stimulation (*6*), heat stress (*7*) and hypoxia (*8*). At the molecular levels, phosphorylation of tyrosine 23 (Y23) of the N-terminal tail of AnxA2 by the Src kinase promotes the cell surface translocation of the (AnxA2•P11)_2_ complex (*7*) while phosphorylation of serine 25 (S25) and/or 11 (S11) by protein kinase C (PKC) negatively impacts the process by disrupting interaction between AnxA2 and P11 (*9*, *10*).

The prototypic second messenger cAMP is a major stress response signal that has been implicated in playing important roles in hemostasis. A recent study shows that cAMP induces the secretion of endothelial von Willebrand factor (VWF), a critical regulator of both primary and secondary hemostasis. The effects of cAMP is further shown to be mediated by a PKA-dependent dephosphorylation of AnxA2 (*11*). However, the role of cAMP in fibrinolysis, especially the effect of exchange protein directly activated by cAMP (EPAC) signaling on AnxA2 translocation, has not been explored and is not clear. In this study, we demonstrate that EPAC1 interacts with AnxA2 and regulates its membrane translocation in endothelial cells. Using genetic and pharmacological approaches, we show that EPAC, acting through the PLCε-PKC pathway, controls (AnxA2•P11)_2_ surface translocation and plasminogen activation. These results suggest that EPAC plays a role in the regulation of fibrinolysis in endothelial cells and may represent a novel therapeutic target for disorders of fibrinolysis.

## EXPERIMENTAL PROCEDURES

*Materials* — Human umbilical vein endothelial cells (HUVECs) and all cell culture reagents were from Lonza (Allendale, NJ). 8-pCPT-2-O-Me-cAMP-AM (007-AM) was from Tocris Bioscience (Bristol, United Kingdom). EPAC inhibitor HJC0758 (>99%) was synthesized and purified as described (*12*, *13*). The following antibodies were purchased from the indicated vendors: mouse monoclonal anti-AnxA2 and anti-S100A10 IgG (BD Transduction Laboratories™, San Jose, CA); rabbit anti-AnxA2 IgG (Santa Cruz Biotechnology, Inc., Dallas, Texas); rabbit anti-pS25-AnxA2 IgG antibody (Thermo Fisher Scientific, Waltham, MA); mouse anti-pY23-AnxA2 IgG (R&D Systems, Minneapolis, MN); rabbit anti-p-PKC (pan) (βII Ser660) IgG, mouse anti-EPAC1 and mouse anti-EPAC2 IgG, HRP-linked anti-rabbit and anti-mouse antibodies (Cell Signaling Technology, Danvers, MA); rabbit anti-PLCε IgG was a gift from Dr. Smrcka’s lab. DyLight 594 Goat Anti-Rabbit IgG antibody and Hard Set Antifade Mounting Medium with DAPI were from Vector Laboratories (Burlingame, CA). Lipofectamine^®^ 2000 was from Thermo Fisher Scientific (Waltham, MA). Plasminogen (Human Plasma) was from Athens Research & Technology (Athens, GA). Chromogenic substrate for plasmin (H-D-Val-Leu-Lys-pNA. 2HCl) was from Molecular Innovations, Inc. (Novi, MI).

*Affinity purification of cellular EPAC1 binding partners* — Hela cells stably expressing FLAG-HA-EPAC1 were cultured in in Dulbecco’s modified Eagle’s medium supplemented with 10% fetal bovine serum. Three confluent 15-cm cultures were lysed in a buffer containing 50 mM Tris (pH 7.5), 150 mM NaCl, 10 mM MgCl2, EGTA 0.5mM, CaCl2 0.5mM, 1% NP-40, and protease and phosphatase inhibitors. The cell lysates were incubated with Anti-FLAG M2 affinity gel overnight at 4 °C with constant gentle mixing, and then washed 5 times in lysis buffer plus 10% glycerol. Bound proteins were eluted with the FLAG peptide. The eluted proteins were then incubated with Anti-HA agarose affinity gel overnight at 4 °C. The affinity beads were washed extensively in lysis buffer plus 10% glycerol and bound proteins were eluted with the HA peptide. The eluate was subject to SDS-PAGE separation and silver staining. Individual protein bands were excised from the gel and subjected to tryptic digestion and peptide mass fingerprinting as described previously (*14*).

*Cell culture—*HUVECs (Lonza) were maintained and sub-cultured in EGM-2™ Endothelial Cell Growth Medium at 37°C in a 5% CO2 humidified incubator. Cells from passages between 2 and 8 were used for experiments described in this study.

*Elution and quantification of cell surface AnxA2* — The levels of cell surface AnxA2 were assessed using an established protocol of cell surface elution with the calcium-chelating agent EGTA as described (*15*). Briefly, confluent HUVECs in 12 well plates were washed once with Hank's Buffered Salt Solution (HBSS) and starved in serum-free EBM (Endothelial Basal Medium) for 2 h prior to the application of external stimuli. After various treatments, the cells were immediately equilibrated to 4 °C on ice for 5 min, then rinsed with ice-cold HBSS, and treated with 10 mM EGTA in HBSS (100 μl/well). Eluates were collected after centrifugation at 500g for 5 min, and analyzed by SDS-PAGE/Western blotting with anti-AnxA2 antibody.

*Immunoblotting analysis* — Total protein lysates from cultured HUVEC cells with various treatments were resolved on 10% or 15% Mini-PROTEAN^®^ TGX Stain-Free™ Precast Gels (Bio-Rad). After electrophoresis, images were taken by ChemiDoc™ Touch Imaging System (Bio-Rad) for total protein loading quantification before proteins were transferred to PVDF membrane (Millipore). The blots were incubated with primary antibodies against AnxA2, p-PKC (pan) (βII Ser660), pS25-AnxA2, pY23-AnxA2, PKCα, EPAC1 and PLCε antibodies at 4°C for overnight followed by incubation with horseradish peroxidase-conjugated secondary antibodies (Cell Signaling Technology, Danvers, MA) and detection using Amersham™ ECL™ Prime Western Blotting Detection Reagent (GE Healthcare Life Sciences, Pittsburgh, PA). The density of autoradiographic signals was assessed with ChemiDoc™ Touch Imaging System (Bio-Rad) and quantitated with Image Lab™ Software (Bio-Rad) or ImageJ software provided by the National Institutes of Health. Protein levels were normalized against an internal control GAPDH or α-tubulin. Eluate AnxA2 protein levels were normalized against the total cellular protein levels. The sample readout was determined as a ratio by dividing the normalized protein level in the treatment cells with that in the control cells, which was set to be 1. Statistical analysis was performed using from at least three independent experiments.

*Immunofluorescence staining of AnxA2* — HUVECs plated on glass coverslips coated with 2% gelatin were treated with 007-AM (5 μM) or vehicle control for 30 min and washed with PBS. For surface AnxA2 staining, the cells were fixed with 2% paraformaldehyde for 10 min, rinse three times in PBS for 5 min each. Fixed cells were incubated with 5% normal goat serum in PBS for 30 min to block non-specific binding and followed by incubation of anti-AnxA2 (1:200; Santa Cruz Biotechnology, Inc.) at 4 °C for overnight. For detecting total cellular AnxA2, cells were fixed in methanol at −20 °C for 20 min, permeabilized with 0.25% Triton X-100 for 5 min and then washed with PBS for three times before the blocking and primary antibody incubation steps. After washing with TBST with 0.1% Tween 20 for three times, cell specimens were incubated in DyLight 594 Goat Anti-Rabbit IgG antibody (1:200) (Vector Laboratories, Burlingame, CA) for 30 min at room temperature in dark, and were mounted with VECTASHIELD Hard Set Antifade Mounting Medium with DAPI after washing with TBST. Fluorescence images were captured using an Olympus microscope (BX51, U-LH100HG) equipped with a Hamamatsu C4742-95 Digital camera. The fluorescence intensity was quantified using the ImageJ program (National Institutes of Health). Five randomly scanned fields of three coverslips from each treatment were analyzed.

*siRNA gene silencing* — Gene-specific Stealth RNAi^TM^ siRNAs oligonucleotides were purchased from Thermo Fisher Scientific and were used at final concentrations of 50 nM for human EPAC1 and PLCε silencing. Stealth RNAi™ siRNA Negative Control, Med GC (Thermo Fisher Scientific) were used as siRNA control. HUVECs seeded on 12-well plates were transfected at 70% confluence using Lipofectamine^®^ 2000 according to the manufacturer’s instructions. After 48 h, cells were starved in serum-free EBM for 2 h followed by treatment of 5 μM 007-AM or vehicle for 30 min. Cells eluates by HBSS/EGTA wash and whole cell lysate were harvested. The effects of siRNAs on gene expression and EPAC-mediated AnxA2 translocation were assessed by immunoblotting analysis.

*Plasmin activity assay* — Cell culture supernatants were assayed for plasmin activity using a chromogenic substrate for plasmin, H-D-Val-Leu-Lys-pNA. 2HCl. HUVECs were grown in 12-well culture plates to confluence. After starvation for 30 min in EBM, cells were treated with 007-AM (5 μM) or HJC0758 (3 μM) for 4 h in the presence of plasminogen at final concentration of 15 mg/ml. Cell culture supernatants were collected, centrifuged at 500 g for 5 min, and assayed for plasmin activity as previously reported (*16*). The final concentration of the substrate was 0.36 mM. The formation of the cleaved product was determined by monitoring absorbance at 405 nm after supernatant sample was incubated with the substrate for 15 min at room temperature. The effect of EPAC1 siRNA on plasminogen activation was also evaluated in a similar manner by measuring the plasmin activity from cell culture supernatants after transfection of EPAC1-specific or control siRNA for 48 h. The average activity of plasmin in the treatment group was normalized to that of the control or vehicle group as fold-change.

*Statistical analyses* — All data were analyzed using the Student’s t-test or the analysis of variance with Bonferroni correction for multiple comparisons between groups. Measurements are expressed as means ± SEM. Statistical significance was designated as P< 0.05.

## RESULTS

### EPAC1 interacts with AnxA2

To identify potential EPAC1 cellular binding partners, we performed tandem immunoaffinity purification of cellular EPAC1-containing protein complexes from Hela cells stably expressing full-length EPAC1 with FLAG and HA epitopes tagged at its C-terminus using immobilized anti-FLAG and anti-HA antibodies as described previously (*14*). A prominent protein band around 36 kD was present in the immunoprecipitated protein complexes sequentially eluted from anti-FLAG and anti-HA matrixes using purified FLAG and HA peptides but missing from the mock purification from Hela cells expressing the control vector only (**Figure 1A**). Tryptic in-gel digestion and peptide mass fingerprinting further revealed the identity of the protein as the AnxA2, which was subsequently validated by immunoblotting using antibody specific against AnxA2 (**Figure 1B**).

**Figure 1.**
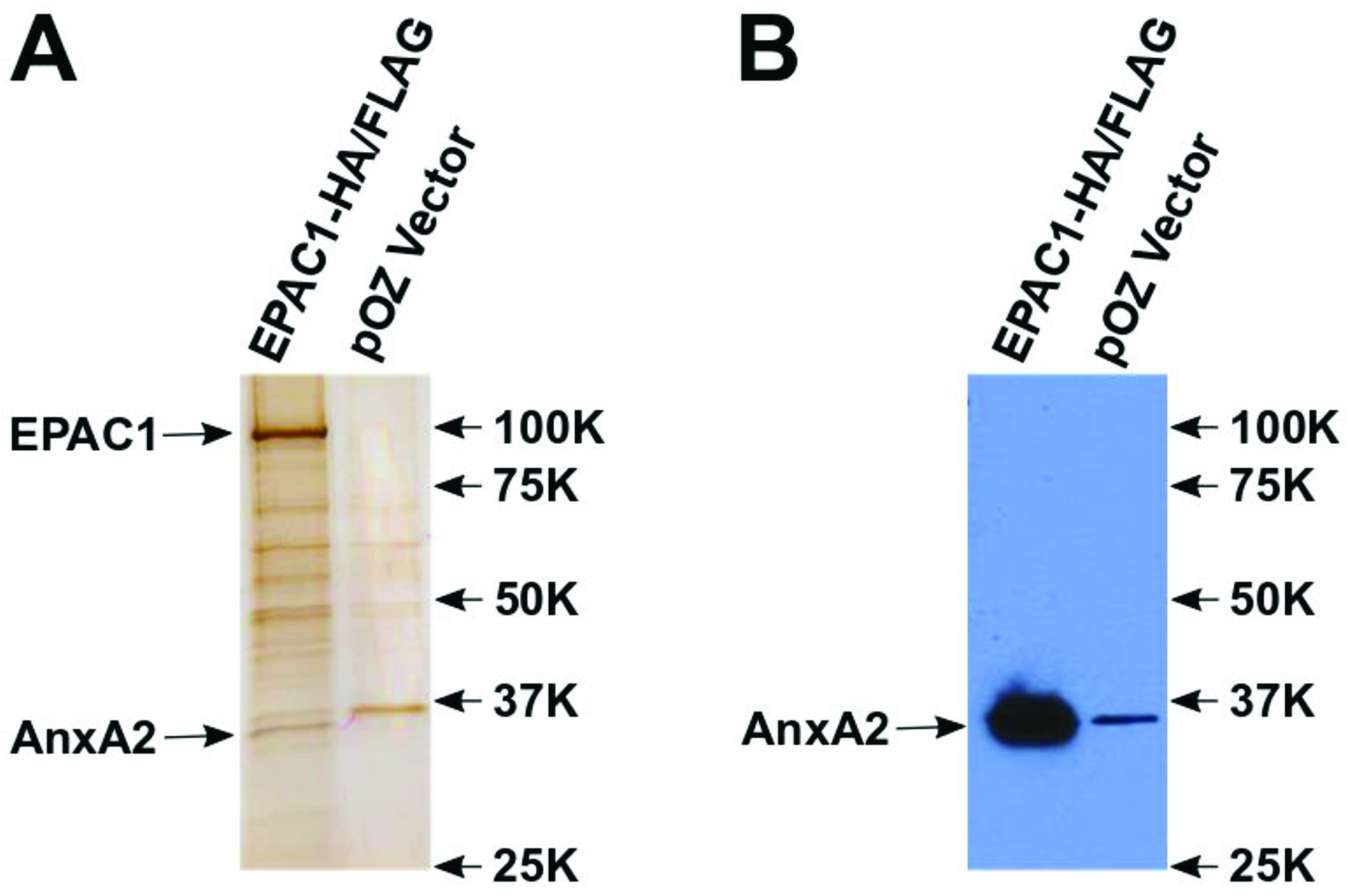
EPAC1 interacts with AnxA2. (A). Silver stained image of cellular protein partners of EPAC1 purified by tandem immunoprecipitation with antibodies specific for FLAG and HA epitopes from Hela cells expressing EPAC1-FLAG-HA fusion protein and control mock purification using vector transfected Hela cells. (B). Affinity pull-down samples from Hela cells expressing EPAC1-FLAG-HA fusion protein or control vector probed with anti-AnxA2 antibody.

### EPAC specific agonist decreases cell surface AnxA2

To explore the functional significance of EPAC1 and AnxA2 interaction, we selectively activated EPAC1 using a membrane permeable EPAC-specific cAMP analog, 007-AM, in HUVECs where both EPAC1 and AnxA2 are expressed abundantly. As shown in **Figure 2A**, selective activation of EPAC1 with 5 μM 007-AM time-dependently reduced the amount of AnxA2 eluted from cell surface by 10 mM EGTA. The decreased cell surface presentation of AnxA2 in response to 007-AM treatment was further confirmed by immunostaining of cell surface AnxA2 of un-permeabilized HUVEC cell (**Figure 2B**). On the other hand, 007-AM treatment did not affect total cellular AnxA2 levels.

**Figure2.**
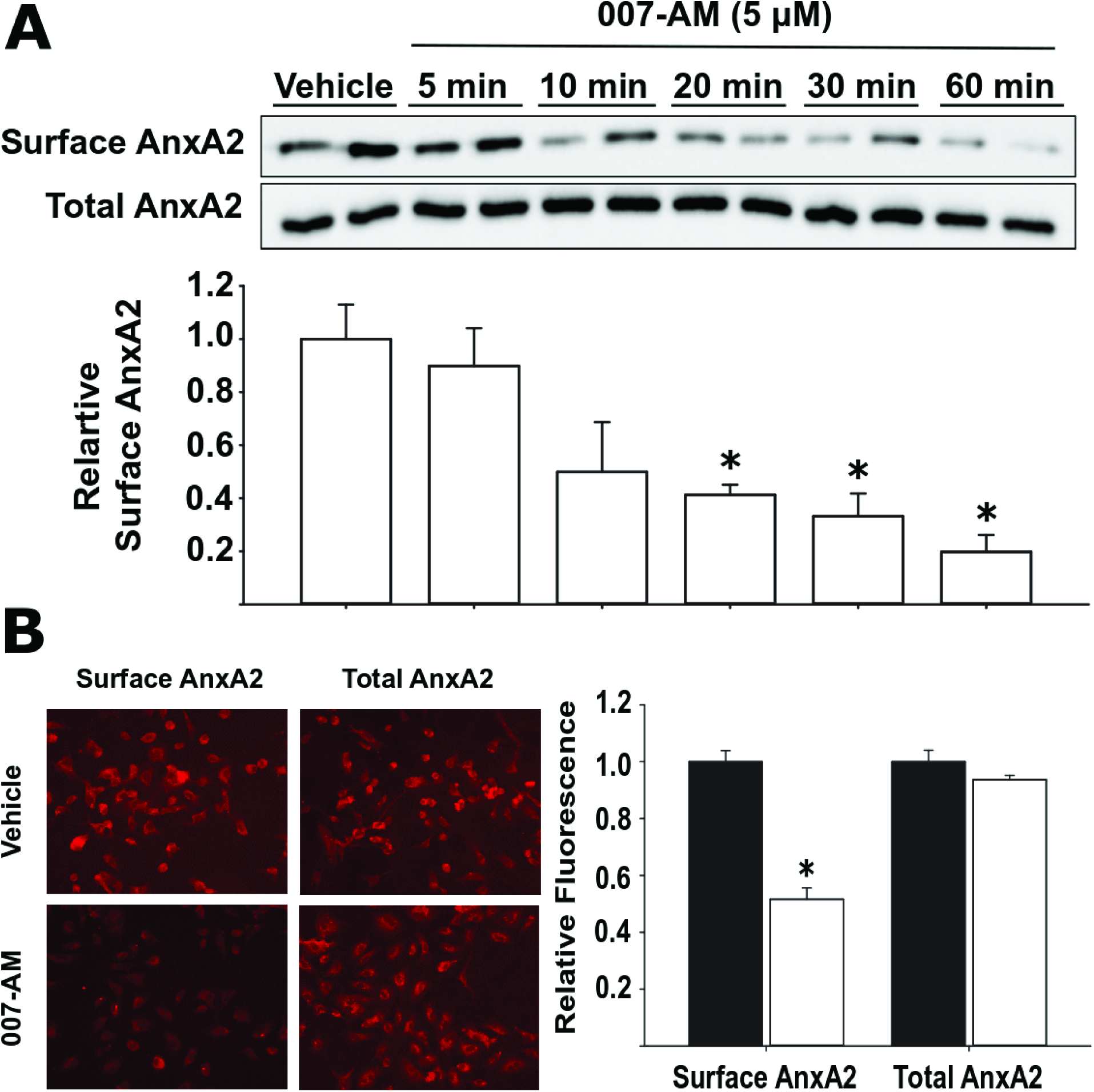
Activation of EPAC by 007-AM time-dependently decreases cell surface presentation of AnxA2. (A) Representative immunoblotting images of cell surface AnxA2 eluted from HUVEC cells treated with vehicle control or 5 μM 007-AM for 5, 10, 20, 30, 60 min. Bar graphs show the relative changes in AnxA2 signals normalized against total whole cell lysate protein loaded in each corresponding lane. (B) Representative immunofluorescence staining images of surface and cytoplasmic AnxA2 in HUVECs treated with 007-AM (5 μM) or vehicle control for 30 min. Bar graphs show the relative immunofluorescence intensity of surface and cytoplasmic AnxA2. Values are expressed as relative fold change in the format of means ± SEM (n = 3 independent experiments). *, P < 0.05 vs vehicle.

### EPAC regulates AnxA2 cell surface translocation by modulating phosphorylation at serine 25 but not tyrosine 23

AnxA2 cell surface translocation is known to be regulated by posttranslational modification events. In particular, phosphorylation of Y23 by Src kinase promotes trafficking of AnxA2 to cell surface while PKC phosphorylation at S11 and S25 suppresses AnxA2 cell surface translocation (*7*). To determine if the effect of EPAC activation on AnxA2 translocation is associated with changes in AnxA2 phosphorylation status, we examined phospho-Y23 and phospho-S25 levels of AnxA2, as well as the level of phospho-PKC, in response to 007-AM treatment. As shown in **Figure 3**, 007-AM induced decrease in cell surface AnxA2 was accompanied by a concomitant increase in phospho-S25 and phospho-PKC levels while the levels of phospho-Y23 remained unchanged. These observations suggest that EPAC-related AnxA2 translocation is not mediated by Src phosphorylation of AnxA2 at Y23, but most likely by PKC phosphorylation at S25.

**Figure 3.**
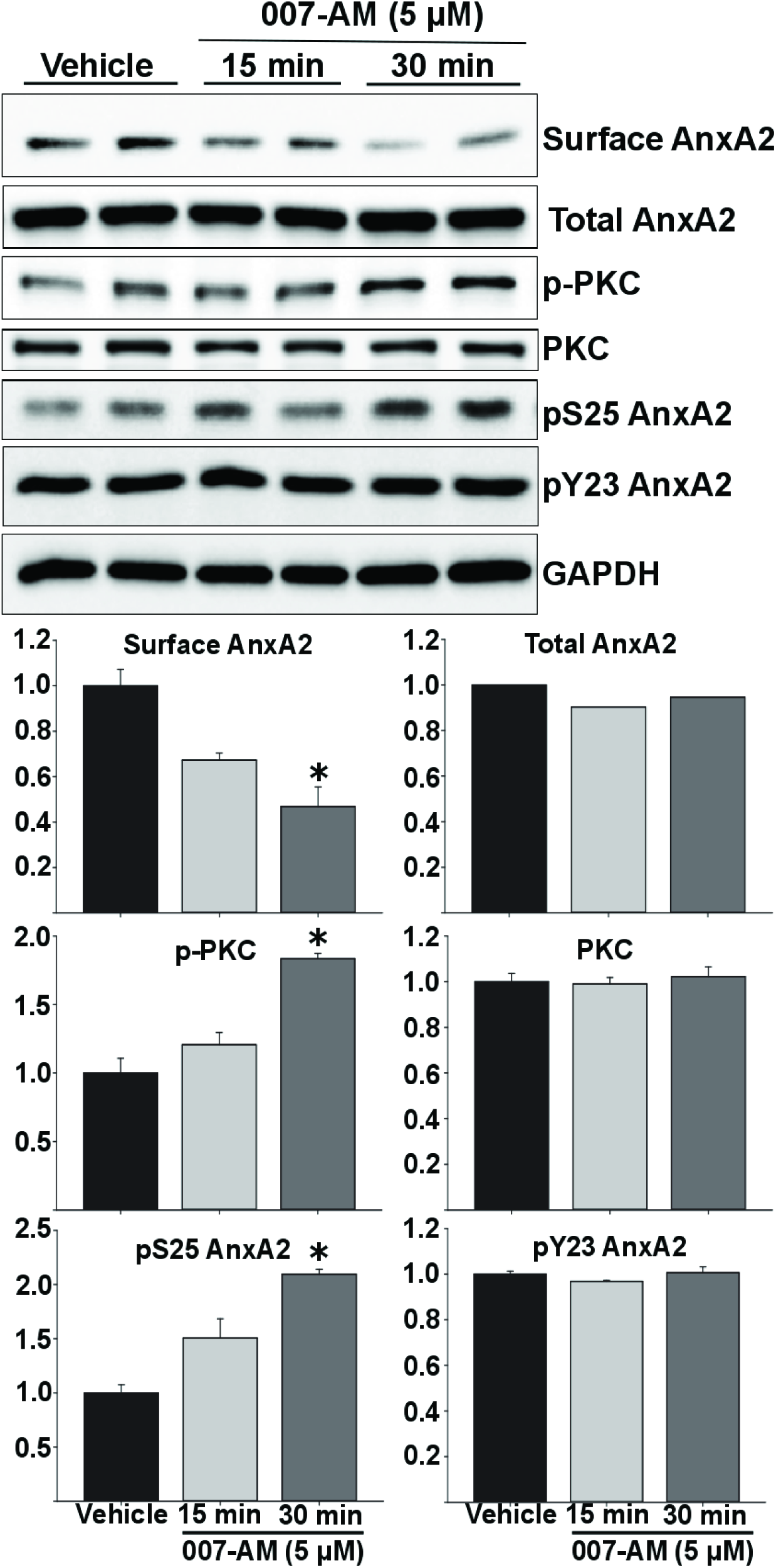
EPAC regulates AnxA2 cell surface translocation by modulating phosphorylation at serine 25 but not tyrosine 23. Representative immunoblotting images of cell surface AnxA2, total AnxA2, p-PKC (pan) (βII Ser660), pS25-AnxA2 and pY23-AnxA2 levels from HUVEC cells treated with vehicle control or in 007-AM (5 μM) for 5 or 30 min. Bar graphs show the relative changes in signal in response to 007-AM stimulation. Values are expressed as relative fold change in the format of means ± SEM (n = 3 independent experiments). *, P < 0.05 vs vehicle.

### Pharmacological inhibition of EPAC enhances AnxA2 cell surface translocation

To further confirm that EPAC activation is responsible for the apparent decreased levels of cell surface AnxA2, we pretreated the HUVEC cells with HJC0758, an EPAC-specific inhibitor, before 007-AM stimulation. As shown in **Figure 4**, HJC0758 treatment reversed the reduction of cell surface AnxA2 and increases in phospho-S25 and phospho-PKC levels induced by 007-AM. In contrast, HJC0758 treatment alone or in combination with 007-AM had no effect on the total cellular AnxA2 level and its phosphor-Y23 level.

**Figure 4:**
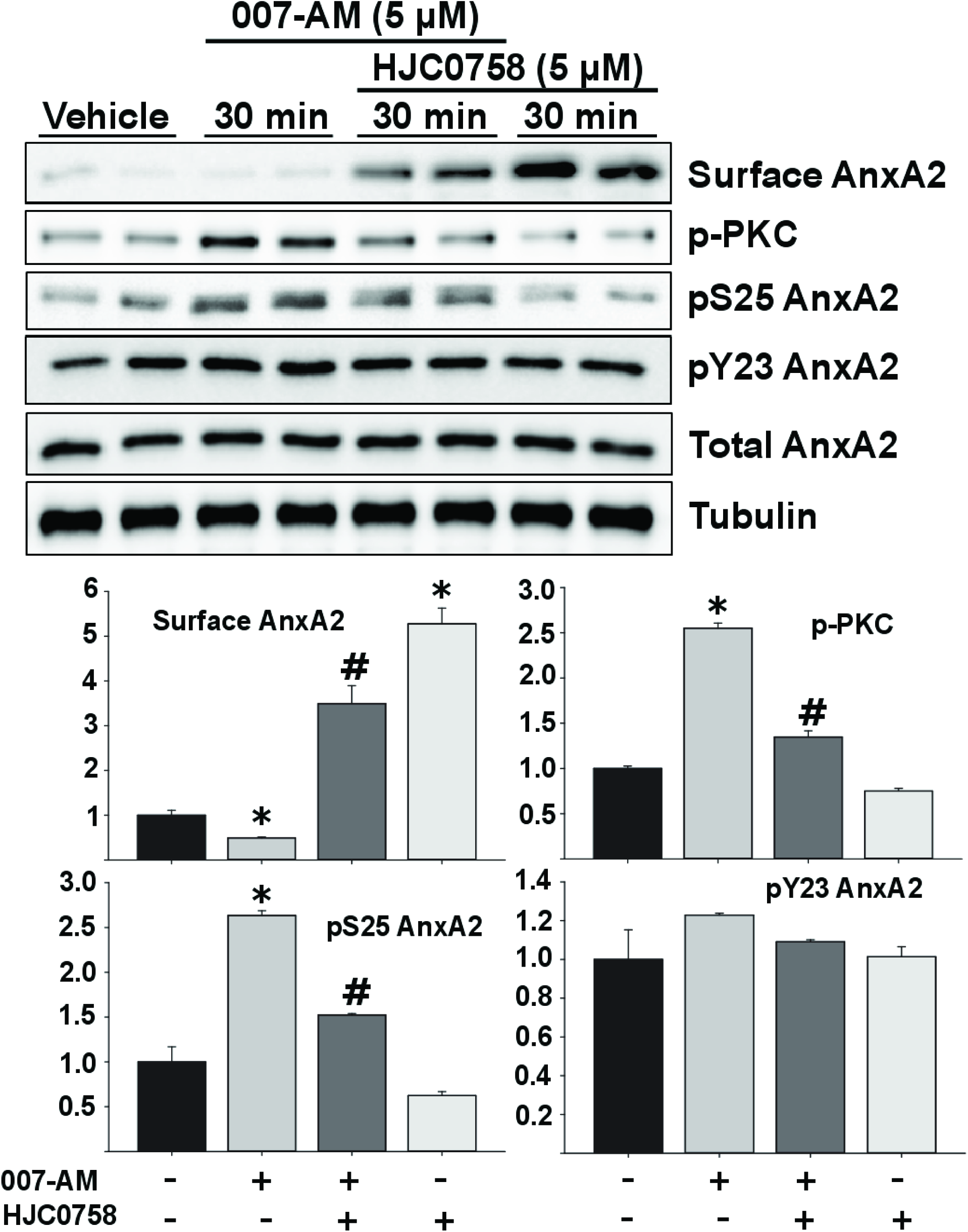
Inhibition of EPAC by HJC-0758 increases surface presentation of AnxA2. Representative immunoblotting images of cell surface AnxA2, total AnxA2, p-PKC (pan) (βII Ser660), pS25-AnxA2 and pY23-AnxA2 levels from HUVEC cells treated with vehicle control or in 007-AM (5 μM) for 30 min in the presence or absence of 30 min preincubation of EPAC specific inhibitor, HJC0758 (5 μM). Bar graphs show the relative changes in signal in response to 007-AM and/or HJC0758 treatments. Values are expressed as relative fold change in the format of means ± SEM (n = 3 independent experiments). *, *P* < 0.05 vs vehicle; *#, P* < 0.05 vs 007-AM only treatment.

### Silencing EPAC1 blocks EPAC agonist-mediated AnxA2 translocation and increases cell surface AnxA2

Between the two EPAC isoform proteins EPAC1, but not EPAC2, is abundantly expressed HUVEC cells (*17*). Indeed, suppressing EPAC1 expression by gene silencing using siRNA led to increased cell surface AnxA2 levels, decreased phospho-S25 and phospho-PKC levels in the presence or absence of EPAC agonist 007-AM (**Fig. 5**). On the other hand, knocking down EPAC1 expression did not alter the total cellular AnxA2 level and its phosphor-Y23 level.

**Figure 5:**
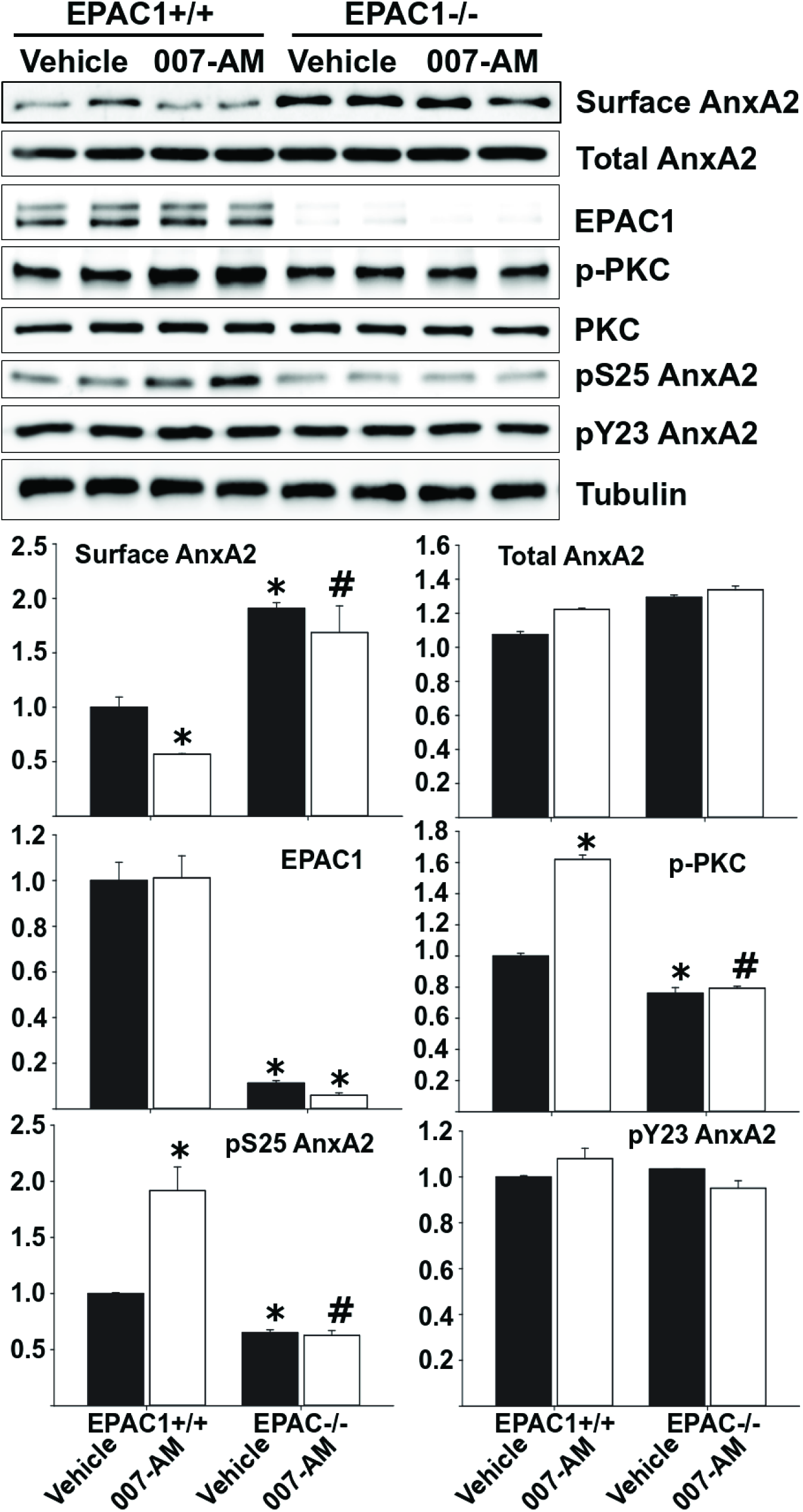
Silencing EPAC1 by siRNA increases surface presentation of AnxA2 in the presence or absence of 007-AM. Representative immunoblotting images of cell surface AnxA2, total AnxA2, p-PKC (pan) (βII Ser660), pS25-AnxA2 and pY23-AnxA2 levels in HUVECs after silencing of EPAC1 by stealth siRNA with or without treatment of 007-AM (5 μM, 30 min). Bar graphs show the relative changes in signal in response to EPAC1 silencing. Values are expressed as relative fold change in the format of means ± SEM (n = 3 independent experiments). *, *P <* 0.05 vs EPAC1^+/+^ vehicle; #, P < 0.05 vs EPAC1^+/+^ 007-AM treatment.

### Silencing PLCε blocks EPAC1-mediated AnxA2 translocation

To determine if EPAC1 acts through phospholipase Cε (PLCε) to regulate AnxA2 membrane translocation and S25 phosphorylation, we examined the effects of suppressing PLCε expression on AnxA2 translocation and phosphorylation by gene silencing using siRNA. As shown in **Figure 6**, knocking down PLCε led to an increase in cell surface AnxA2 levels and a concomitant decrease in phospho-S25 and phospho-PKC levels while the total cellular AnxA2 level and its phosphor-Y23 level remained unchanged. More importantly, down-regulation of PLCε completely blocked 007-AM induced inhibition of AnxA2 cell surface translocation, as well as 007-AM induced activation of S25 and PKC phosphorylation. In fact, the level of cell surface AnxA2 remained elevated while the levels of phospho-S25 and phospho-PKC levels stayed repressed in PLCε knockdown HUVEC cells in the presence of EPAC stimulating agent, 007-AM. These observations are indicative that PLCε/PKC signaling acts down-stream of EPAC in mediating cAMP’s effect on modulating AnxA2 functions.

**Figure 6:**
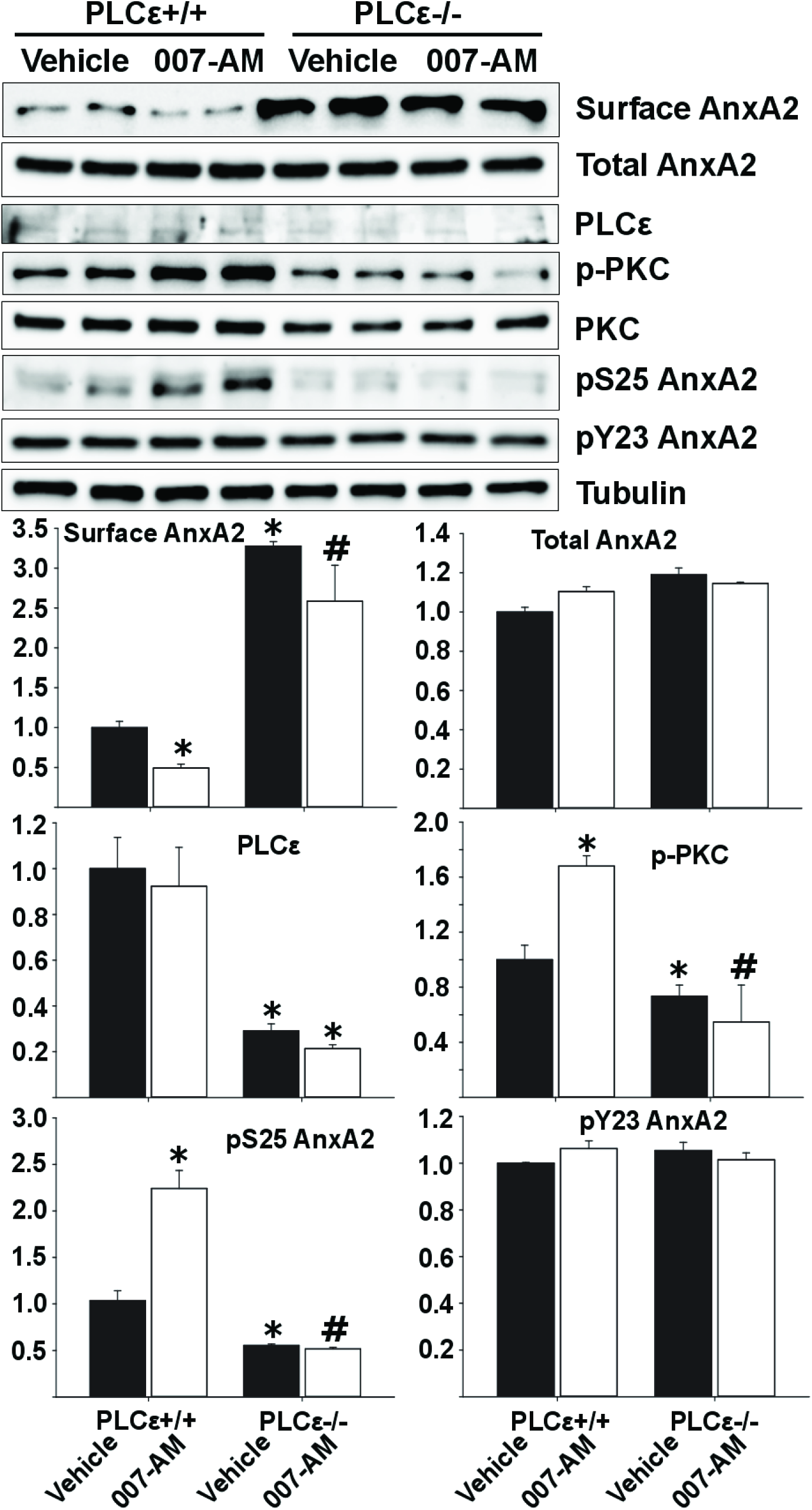
Silencing PLCε by siRNA increases surface presentation of AnxA2 in the presence or absence of 007-AM. Representative immunoblotting images of cell surface AnxA2, total AnxA2, p-PKC (pan) (βII Ser660), pS25-AnxA2 and pY23-AnxA2 levels in HUVECs after silencing of PLCε by stealth siRNA with or without 007-AM (5 μM, 30 min) stimulation. Bar graphs show the relative changes in signal in response to PLCε silencing and/or 007-AM treatment. Values are expressed as relative fold change in the format of means ± SEM (n = 3 independent experiments). *, *P* < 0.05 vs PLCε^+/+^ vehicle; #, P < 0.05 vs PLCε^+/+^ 007-AM treatment.

### EPAC regulates plasminogen activation

As an endothelial co-receptor for plasminogen and tissue plasminogen activator, the cell surface AnxA2/P11 tetramer complex promotes plasminogen activation (*18*). To examine if EPAC is capable of regulating plasmin generation through modulating AnxA2 translocation in endothelial cells, we monitored the activities of conversion of plasminogen to plasmin in HUVEC cells in response to EPAC activation and inhibition. As shown in **Figure 7A**, stimulating HUVEC cells with 007-AM led to a significant decrease in plasmin activity while inhibition of EPAC1, pharmacologically by an EPAC specific inhibitor HJC0758 or genetically by EPAC1 siRNA, promoted the formation of plasmin (**Fig. 7B & C**).

**Fig. 7.**
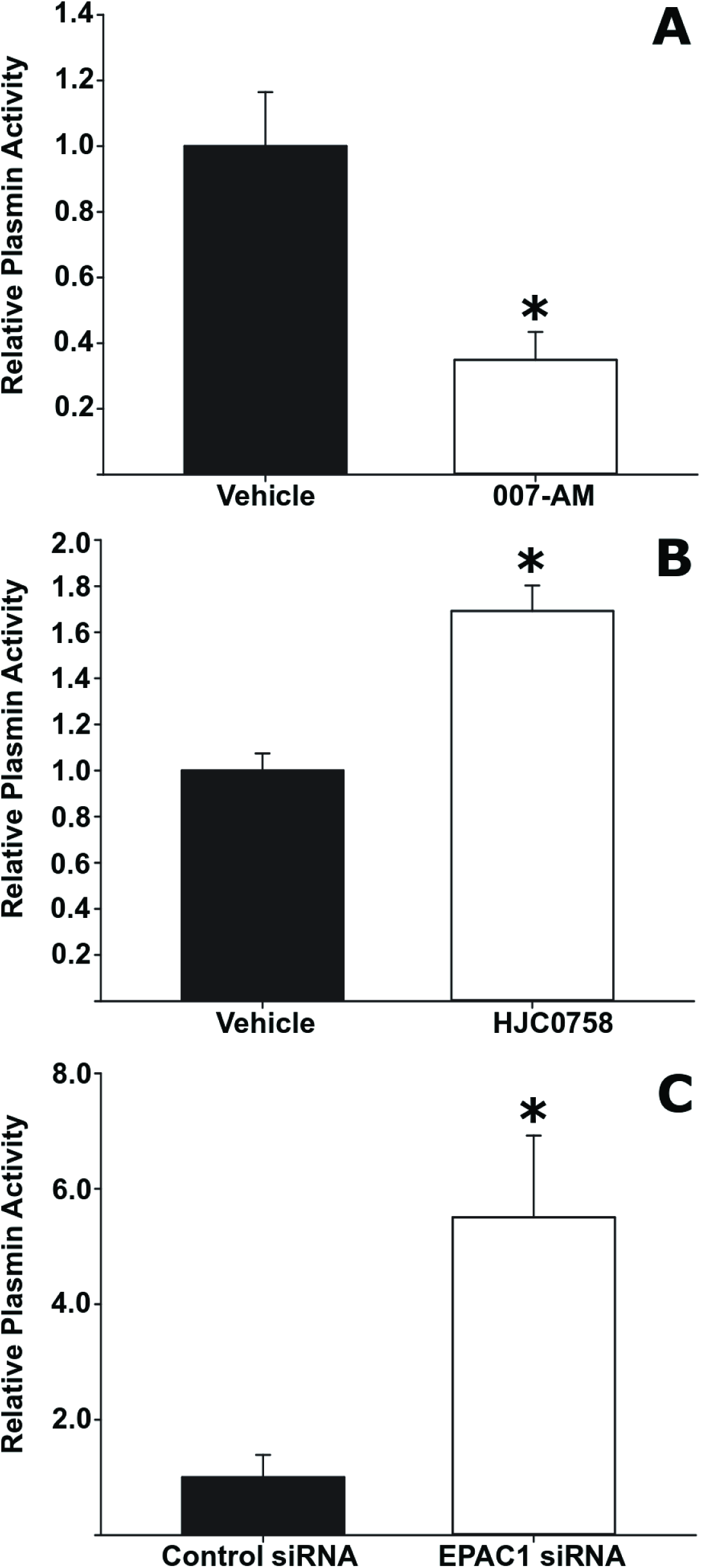
The effect of EPAC1 activation or inhibition on plasminogen conversion into plasmin. Plasmin activity generated from plasminogen activation by culture medium collected from HUVEC cells treated with 007-AM (5 μM) or vehicle for 30 min (A); or with HJC0758 (3 μM) or vehicle for 30 min (B), or from HUVEC cells transfected with EPAC1 or control siRNA (C). Values are expressed as relative fold change in the format of means ± SEM (n = 3 independent experiments). *, P < 0.05 vs vehicle (A & B) or control siRNA (C).

## DISCUSSIONS

cAMP is a key stress-response signal that has been implicated in regulating major endothelial cell functions such as cell permeability, junction stability, inflammatory response and hemostasis (*19*, *20*). cAMP exerts its effects mainly through the activation of protein kinase A (PKA) and EPAC (*21*). While the roles of EPAC signaling in modulating endothelial barrier functions and inflammatory response have been examined extensively (*22*, *23*), studies and information related to cAMP’s functions in fibrinolysis are scarce. Our current study reveals that EPAC1 interacts with AnxA2, a phospholipid binding protein abundantly expressed in endothelial cells and important for vascular fibrinolysis and integrity (*24*, *25*). This unexpected finding led us to investigate the physiological functions of EPAC1:AnxA2 interaction. Using genetic and pharmacological approaches, we show that activation of EPAC1 suppresses AnxA2 cell surface presentation in endothelial cells. Stimulation of HUVEC cells with a membrane permeable EPAC-specific agonist 007-AM (*26*), results in a time-dependent decreases in AnxA2 cell surface presentation as measured by immunoblotting of cell surface EGTA elution, an established protocol for quantifying cell surface AnxA2 (*15*), as well by direct visualization of immunostaining of cell surface AnxA2. Conversely, inhibition of EPAC1 by an EPAC specific antagonist HJC0758, a second generation EPAC inhibitor based on lead compound ESI-09 (*13*, *27*), or by EPAC1-specific siRNA promotes the surface translocation of AnxA2. Taken together, these observations provide convincing evidence to support a role of EPAC1 in regulating AnxA2 cellular trafficking in response to various stress signals that act through the cAMP signaling axis.

AnxA2 was initially identified as a major cellular substrate of oncogenic Src kinase of Rous sarcoma virus (*28*). Subsequent studies revealed that Src kinase phosphorylated AnxA2 at residue Y23 (*29*, *30*) and phosphorylation of Y23 is required for stress-induced cell surface translocation of (AnxA2•P11)_2_ tetramer complex (*7*). In addition to tyrosine phosphorylation, AnxA2 is also known to undergo serine/threonine phosphorylation by PKC (*31*, *32*). While AnxA2 translocation to cell surface is positively regulated by Src tyrosine phosphorylation, PKC phosphorylation inhibits AnxA2 membrane translocation (*10*). Two PKC phosphorylation sites, S11 and S25, located within the N-terminal tail of AnxA2 have been identified: serine 25 was reported to be phosphorylated by PKC both *in vivo* and *in vitro* (*32*), while serine 11 was only observed to be phosphorylated *in vitro* (*33*, *34*). Moreover, S25 phosphorylation of AnxA2 was associated with the nuclear entry of AnxA2, DNA synthesis and cell proliferation, whereas serine 11 has no obvious influence (*35*). These reports support the notion that S25 is the major physiological PKC phosphorylation site of AnxA2. Using phosphor-specific antibodies for pY23-and pS25-AnxA2, we show that alteration in cell surface presentation of AnxA2 in response to EPAC1 activation or inhibition is accompanied by reciprocal changes in pS25-AnxA2 and p-PKC levels while the level of pY23-AnxA2 remains constant. These results suggest that EPAC signaling modulates AnxA2 cell surface translocation via promoting S25 phosphorylation through PKC activation. Since the initial pivotal discovery by Schmidt and colleagues (*36*), it is now well-established that EPAC signaling can act through PLCε to activate PKC. Our data show that silencing PLCε by siRNA not only enhances AnxA2 cell surface presentation but also blocks EPAC’s ability to suppress AnxA2 membrane trafficking in response to stimulation by 007-AM, an EPAC-specific agonist, demonstrating that PLCε functions as a down-stream effector of EPAC1 signaling in modulating AnxA2 translocation.

As a profibrinolytic co-receptor for plasminogen and tPA, (AnxA2•P11)_2_ complex is capable of stimulating the activation of the major fibrinolysin, plasmin at endothelial cell surfaces (*37*). Mice deficient in AnxA2 display increased fibrin accumulation within blood vessels and impaired clearance of injury-induced thrombi (*24*). Anti-AnxA2 auto-antibodies have been reported in patients with thrombosis-associated diseases, such as antiphospholipid syndrome, pre-eclampsia and cerebral venous thrombosis (*38*). On the other hand, overexpression of AnxA2 in acute promyelocytic leukemia patients leads to a bleeding diathesis due to excessive cell surface AnxA2-dependent generation of plasmin (*39*). The ability of EPAC1 signaling to modulate endothelial AnxA2 cell surface presentation suggests a potential role of EPAC1 in regulating vascular fibrinolysis. Indeed, our study shows that manipulating EPAC1 activation in human endothelial cells leads to corresponding changes in plasmin activation at cell surfaces. Although additional studies using animal models and/or clinically related samples are required to further validate EPAC1’s functions in hemostasis and fibrinolysis. Our findings, nonetheless, suggest a potential therapeutic strategy for treating disorders associated with vascular fibrinolysis by targeting EPAC1/ PLCε signaling.

## Acknowledgments

This work was supported by National Institutes of Health grant R01GM066170, R35GM122536 and R01AI111464.The funders had no role in the study design, data collection and analysis, decision to publish, or preparation of the manuscript. We thank Ms. Pei Luo for helping designing the real-time PCR primers. We are indebted to Dr. Alan V. Smrcka’s lab (University of Michigan) for their generous gifts of anti-PLCε antibody and other related reagents for this study.

## Author contributions

WY: designing research studies, conducting experiments, acquiring and analyzing data and manuscript preparation; FCM: designing research studies, conducting experiments, acquiring and analyzing data. XC: designing research studies, acquiring and analyzing data and writing the manuscript.

## Additional Information

Competing financial interests: The authors declare no competing financial interests.

